# Genome assembly and diagnostic DNA markers for sex of the largest freshwater fish in North America, the white sturgeon (*Acipenser transmontanus*)

**DOI:** 10.1101/2025.11.14.688503

**Authors:** Stuart C. Willis, Jeremiah Smith, Shawn R. Narum

**Author notes:** Corresponding author’s.

## Abstract

Sturgeon and paddlefish represent some of the most early diverging branches of ray-finned fishes and have undergone at least one global and several lineage-specific whole genome duplication events. White sturgeon (*Acipenser transmontanus*), the largest freshwater fish in North America, have experienced at least two rounds of whole genome duplication, and may exhibit both di- and multi-valent meiotic segregation. Moreover, they exhibit contemporary ploidy variants due to spontaneous autopolyploidy, particularly in aquaculture. Nonetheless, as a species with several population segments that exhibit chronic recruitment failure, conservation aquaculture is an important part of their management. To facilitate the development of genetic tools to aid white sturgeon conservation, as well as a basis to understand how ploidy changes and variation historically and contemporarily shape the evolution of this species, we present a genome assembly for a white sturgeon from the Snake River, Idaho, USA. Analysis of sequence data used for assembly indicated a haploid genome size of approximately 1.5Gbp, implying tetraploidy (4N), while analysis of heterozygous k-mers from 21 to 41 bp suggest the genome reflects both 4N and 8N variants. The final genome assembly, scaffolded using linkage maps constructed from Fraser River and Snake River F1 families, contained 6.26Gbp in 832,145 scaffolds, consistent with published genome size estimates. Conserved ortholog completeness for this genome (90.5%; 22.8% single-copy and 67.7% duplicated) was similar to the putatively diploid sterlet sturgeon, and the largest linkage map-based scaffold was 55.2Gbp, though the N50 for this assembly was only 416Kbp, indicating the assembly remains fragmented. We demonstrate the utility of this assembly by identifying genomic regions significantly associated with sex. Genetic markers, designed for inclusion in an amplicon genotyping panel, predicted sex 96.6% and 81.5% correctly in females and males, respectively, providing a strong overall association (□^2^ p-value < 2.7×10^-37^) with some variation by geographic region.

**Article summary:** Functional polysomes (more than two chromosomes that pair in meiosis) are rare among vertebrates. White sturgeon, the largest freshwater fish in North America, are tetraploid and occasionally hexaploid. Several populations of this species are stagnant or declining, requiring aquaculture to bolster reproduction. We analyzed whole genomic data and assembled the genome of an individual from the unique Snake River, Idaho, population. Results indicate most genetic variants are consistent with tetraploidy, and the genome assembly, while fragmented, is largely complete. We demonstrate its utility by designing genetic markers for sex for use in conservation and commercial aquaculture.

## Introduction

Sturgeons and paddlefishes, fishes in the order Acipenseriformes, are often referred to as ‘living fossils’. While somewhat misleading, this moniker reflects that they were an early-diverging branch within the Actinopterygii or clade of ‘ray-finned’ fishes, have exhibited conservative morphological change relative to ancient acipenseriform fossils, and unlike most other ray-finned fishes, exhibit only a weakly ossified, cartilaginous skeleton (Peng et al., 2007). In contrast, relative to the estimated ancestral Actinoperygiian karyotype, sturgeon and paddlefish have undergone several whole genome duplications (Peng et al., 2007), and while many species appear to have made considerable if not complete evolution back to functional diploidy, some remain polyploid and potentially multivalent across much of the genome (Fontana, 2002). Moreover, in some sturgeon species, spontaneous autopolyploidy results in ploidy variation within a single population, although the origin and long-term demographic importance of these variants is not well-understood (Drauch Schreier et al., 2011). Nonetheless, these observations make sturgeons some of the few contemporarily polyploid, fully sexually reproducing vertebrate species (Jighly et al., 2018; Spoelhof et al., 2017).

Sturgeon and paddlefishes are often known for their size and unique, even bizarre morphology, which makes them a conspicuous target for commercial and recreational fisheries, as well as for being the source of roe for caviar. Most contemporary caviar and sturgeon meat, however, comes from commercial aquaculture operations, since most natural populations were overharvested if not threatened or endangered by the mid-twentieth century, and at least one acipenseriform, the Chinese paddlefish, was declared extinct in the wild in 2019 (Lobanov et al., 2024). Indeed, the ‘periodic’ life history strategy of these fishes makes them extremely vulnerable to overfishing (Kopf et al., 2025; Winemiller, 2005). They exhibit individuals that mature after a decade or more, experience very low rates of natural mortality with adults that live for decades, if not more than a century, and have high fecundity. Notably, however, survivorship among early life stages is low and depends on environmental conditions that may occur unpredictably in time and space, including flow rates and substrate composition (Parsley et al., 1993; Parsley & Beckman, 1994). In addition to overharvest, anthropogenic alteration of natural flow patterns and habitat fragmentation have resulted in species with reduced population sizes that now must also contend with diminished or failed recruitment (Mallette, 2014). As such, there are numerous population management programs for sturgeons and paddlefish across their range, including the development of conservation aquaculture programs to bolster threatened populations (Korman et al., 2025). The success of these aquaculture programs, however, depends on the efficient production of hatchery progeny which reflect the genetic diversity and sex-ratios of natural-origin populations, an endeavor which can be greatly aided by molecular markers and especially markers for sex, notwithstanding the ancestrally and possibly functionally polyploid nature of the genomes.

The white sturgeon (*Acipenser transmontanus*) typifies all of the challenges exhibited by acipenseriform fishes. White sturgeon are the largest ‘freshwater’ fish in North America, historically reaching lengths over 4m. Although they are obligate freshwater spawners, white sturgeon are considered to be quasi-amphidromous and range along the Pacific coast from Baja California to Alaska (Hildebrand et al., 2016). Despite their extensive range, there are only three known self-sustaining populations, in the Sacramento-San Joaquin, Fraser, and Columbia River Basins, with the latter estimated to be the largest (Hildebrand et al., 2016), despite the strong segmentation of the basin by hydropower facilities. Within the Columbia Basin, as elsewhere, white sturgeon populations were severely overharvested into the twentieth century and, despite modern protections, are imperiled by limited recruitment in many areas.

White sturgeon have a karyotype of ∼270 chromosomes (Van Eenennaam et al., 1998), although there is considerable uncertainty of their genome size (C-value), with estimates ranging from 5.1 to 9.6pg (Blacklidge & Bidwell, 1993; Hinegardner, 1976). Relative to the Actinopterygiian median (2N ∼48), this karyotype indicates considerable genome expansion within Acipenseriformes, though contemporary congeners with ∼120 chromosomes are considered functional diploids (Fontana, 2002). Indeed evidence for white sturgeon suggests they exhibit tetravalent or bivalent segregation and inheritance patterns (Delomas et al., 2021; Van Eenennaam et al., 1998), though the proportions of each across the genome are difficult to estimate, and for simplicity they are considered tetraploid (4N) relative to sturgeon species with ∼120 chromosomes (Fontana et al., 2008; Gille et al., 2015). Moreover, spontaneous autopolyploidy is not uncommon in hatchery white sturgeon, and occasionally among natural-origin fish as well, producing hexaploid offspring (6N, relative to 2N≍120), which themselves produce viable pentaploid (5N) offspring when backcrossed (Drauch Schreier et al., 2011; Gille et al., 2015). Notably, female white sturgeon are predicted to be the heterogametic sex (ZW sex determination), like other sturgeons (Fopp-Bayat et al., 2007; Van Eenennaam et al., 1999).

However, if autopolyploids are produced through the retention of the second polar body of the egg, as hypothesized (Gille et al., 2015), meaning maternal genetic contributions are doubled, sex ratios, fertility, and perhaps viability may commonly be compromised among spontaneous autopolyploids and backcross offspring (Wertheim et al., 2013). Despite these potential consequences, ploidy variation clearly has long played a role in sturgeon evolution (Redmond et al., 2023), but deciphering how rediploidization following repeated genome duplications has resulted in both susceptibility and resilience to contemporary ploidy variation in white sturgeon has been limited by genomic resources for it. Here, we present a draft genome assembly for white sturgeon produced from chromatin proximity ligation, microfluidic fragment barcoding, and linkage disequilibrium mapping. We demonstrate the utility of this genomic assembly by identifying regions of the assembly that are homologous to sex-associated regions of other sturgeon and testing the efficacy of molecular markers for them at predicting phenotypic sex in white sturgeon.

## Methods

### Initial genome assembly

Whole blood from an adult female white sturgeon originating from the Snake River population (a tributary to the Columbia River) near Hagerman, Idaho, USA was submitted to NRGene (Ness Ziona, Israel) for paired-end, mate-pair (proximity-ligation) and 10X Chromium (microfluidic fragment barcoding) sequencing (Table 1). To estimate the genome size and ploidy status of the sequenced individual, we utilized the ≤470bp-insert, 2×250bp library with Genomescope and Smudgeplot (Ranallo-Benavidez et al., 2020). For each analysis, kmers were tallied with jellyfish (Marçais & Kingsford, 2011) and FastK (Myers, 2020), respectively, with kmer lengths of 21, 23, 31, and 41 bp. Genomescope analysis was run with assumed ploidy of 2N and 4N (maximum for this analysis is 6N).

**Table 1.**
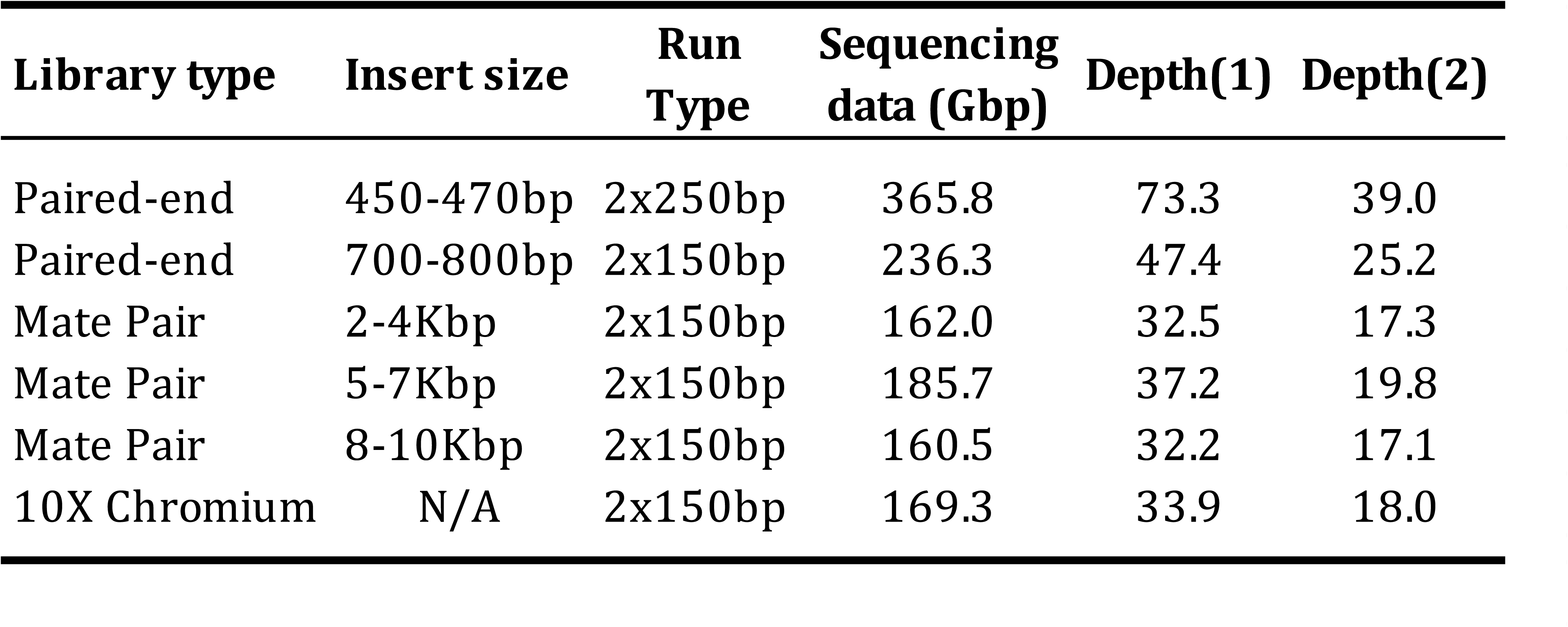
Sequence data statistics. Coverage (depth) is listed for published genome estimates of (1) 5.1pg; (Hinegardner 1976) and (2) 9.6pg (Blacklidge et al. 1993).

Sequence data were utilized with the DeNovoMAGIC v3 assembler to produce a phased (allelic) genome assembly (NRGene). Completeness of the phased assembly was assessed using Compleasm v0.2.7 (Huang & Li, 2023) based on the Actinopertygii ortholog set from orthoDB v12, which contains 7,207 orthologous genes. For comparison, similar evaluations of completeness were made on the NCBI RefSeq (haploid) assembly for the sterlet sturgeon (*Acipenser ruthenus*), a putatively diploid European sturgeon with 2N ∼120 chromosomes (GCF_902713425.1) (Du et al., 2020).

### F1 family sequencing and variant calling

Linkage mapping was made from two outbred F1 families. The first used restriction-site associated DNA sequencing (RADseq) of a mixed reduced- and normal-ploidy family of white sturgeon originating from the Fraser River population (British Columbia, Canada) and the second used messenger RNA sequencing (RNAseq) of a normal-ploidy family from the Snake River population.

For the Fraser River F1 family, ultraviolet light irradiated sperm were used to fertilize eggs from a female white sturgeon at Vancouver Island University (British Columbia, Canada). Whole genomic DNA from somatic (fin) tissue from 226 offspring plus 2 parents was utilized in a restriction digest with a single enzyme with a 8bp recognition sequence (SbfI) following published protocols (Baird et al., 2008; Micheletti et al., 2018). Barcoded libraries were sequenced on an Illumina NextSeq 500 (2×150bp). Reads were mapped to the genome assembly using Hisat2 v2.2.1 (Kim et al., 2019) with the--no-spliced-alignment option enabled, and SNPs were called using bcftools v1.22 (Danecek et al., 2021) jointly using all individuals and prospective parents, then filtered using vcftools v0.1.17 (Danacek et al., 2011) to separate diploid vs haploid mapping families and retain SNPs with minor allele frequencies greater than 0.1 and with missing genotypes in fewer than 50% of segregants within a mapping family. Genotypes in VCF format were converted to joinmap format using the populations pipeline from STACKS v2.68 (Catchen et al., 2011).

For the Snake River F1 family, sampling occurred at the Idaho Springs Foods sturgeon hatchery near Hagerman, ID, USA. Dorsal fin punches were collected from the and preserved in RNAlater. Whole bodies of 92 offspring were collected and preserved in RNAlater and stored at −20C until RNA isolation. Larvae were ∼3cm in length at the time of collection and were euthanized through cervical dislocation. Total RNA was isolated from a mixed-tissue portion of the head of each offspring and fin clips from the parents using the Direct-zol RNA Miniprep Plus Kit using manufacturer protocols (Zymo Research, Irvine, CA, USA). Quality for each individual extract was assessed utilizing High Sensitivity RNA ScreenTape Analysis (Agilent Technologies INC, Clara, CA, USA). Individuals with RNA integrity numbers ≥ 7 were used in downstream library preparation with a total of 92 offspring and 2 parents used. We isolated messenger RNA (mRNA) from the total RNA using Poly(A) mRNA Magnetic Isolation Module (New England Biolabs, MA, USA).

RNA-seq library preparations were performed using half volume reactions with the NEBNext Ultra II RNA Library Prep Kit for Illumina (New England Biolabs, MA, USA), which produces non-stranded cDNA fragments. We used an 8-minute mRNA fragmentation time to ensure fragment sizes were ≥ 350bp prior to adapter ligation and used 13X cycles during PCR enrichment. Bead cleaning was omitted prior to PCR enrichment but two or three bead cleans were performed post-PCR to ensure dimer removal. Equimolar pools of two sets of 46 offspring plus the two parents were created for sequencing on an Illumina NextSeq2000 using a P3 300 cycle kit with 2×150bp chemistry (two libraries of 46 unique offspring; each parent sequenced twice). Data were quality-trimmed with BBmap (Bushnell, 2015) then mapped to the phased assembly using HiSat2 as unstranded cDNA and retaining secondary alignments. Only properly paired alignments were retained and used to score variants with bcftools, which assumed diploidy. Variants were filtered for a minimum quality of 20, minimum depth of 10 reads, maximum missing data of 10%, and minimum allele frequency of 0.1.

Parents and offspring from both F1 families were also genotyped for a panel of 291 anonymous, autosomal SNPs developed for white sturgeon (Delomas et al., 2021). These were amplified and genotyped using GT-seq (Campbell et al., 2015), with a minimum of 20 reads, using sequences from an Illumina NextSeq 500 (1×75bp). These SNP data, as well as the RADseq data, were used to distinguish Fraser River F1 family members that were reduced versus normal ploidy (see Delomas et al., 2021 for details), since not all sperm were effectively irradiated to eliminate the male contribution.

### Linkage map construction and phased assembly scaffolding

Further scaffolding of the phased assembly was performed using genetic linkage maps from segregating polymorphisms within the three sibships described above. Specifically, 1) polymorphisms from RAD-seq data from reduced-ploidy (hereafter ‘haploid’) Fraser River sibships, 2) polymorphisms from RAD-seq data from normal-ploidy (hereafter ‘diploid’) Fraser River sibships, and 3) polymorphisms from RNA-seq data from the normal-ploidy Snake River sibships. For each dataset, we initially constructed a high-confidence linkage map using a subset of markers with highly replicated genotypes (those wherein the same segregation patterns across all offspring were observed 5 or more times).

Linkage analysis was performed for each mapping family separately using JoinMap5 using maximum likelihood mapping under the appropriate crossing design for each sibship: “Haploid” for the haploid sibship and “outbred F1” (CP: cross pollinator) for the other two (Stam, 1993). Additional markers were added to this map if >80% of individuals had identical genotypes to framework markers when considering segregation in both coupling and repulsion phases. We then used these three maps as a framework to further scaffold the initial assembly using Allmaps from JCVI utility libraries 1.4.2 (Tang et al., 2015), with weights for each position scaled to the number of genotype mismatches relative to a framework marker (mismatches/weight: 0/20, 1/8, 2/4, 3/2, 4/1) within each map, initially prioritizing contigs that were captured by the haploid map. Subsequent scaffolding rounds captured additional scaffolds that were absent from the haploid map, prioritizing the diploid map, and then those that were represented only in the RNA-seq-derived map. We evaluated the resulting scaffolds using the geneticmap.py LD function of JCVI.

### Comparison to the sterlet assembly

To assess the large-scale structure of the scaffolded assembly and identify orthologous (sturgeon/sterlet) or paralogous (sturgeon/sturgeon) chromosomes, we aligned our white sturgeon assembly to the sterlet genome assembly (GCF_902713425.1) using minimap v2.28-r1209 (Li, 2016) with the-asm20 option enables for divergence assembly to reference mapping. Alignments were visualized using DGENIES v1.5 (Cabanettes & Klopp, 2018). An alignment of white sturgeon scaffolds to themselves was similarly made.

### Genetic markers for phenotypic sex

In comparison of the published female and male assemblies of sterlet sturgeon (*A. ruthenus*), Kuhl et al. (2021) identified two sex-specific, “unalignable” regions (∼10Kbp for males, ∼16Kbp for females) embedded in an otherwise homologous region, with males showing little to no coverage for the female-specific region (putative W chromosome). We extracted the sex-specific and flanking regions from these assemblies and searched for similar sequences in our phased white sturgeon assembly using NCBI *blastn* (Camacho et al., 2009) with minimum e-value 1×10^-5^ and 50% similarity. This identified three phased scaffolds containing homologous sequences: two putative Z and 1 putative W scaffolds. Alignments made with Mummer v4 (Marçais et al., 2018) (*nucmer--mum--nosimplify -c 100 -g 1000*) confirmed the homology of the flanking regions, as well as the sex-specific regions for the respective scaffolds. From alignments of all of these, made in Geneious (Biomatters), we identified a small (∼5Kbp) island of homology within the otherwise unalignable, sex-specific regions which contained SNP variants distinguishing the putative Z and W scaffolds (named *zwhr*, for ‘Z/W homologous region’). We developed primers for two small amplicons (<100bp), one covering a subset of these SNPs in the nested, homologous region, and the other a putative-W-specific amplicon from the adjacent, unalignable region (named *p455160ex*, for ‘phased scaffold 455160 extraction’). We tested the utility of these markers to predict sex phenotype in known sex sturgeon from the Columbia River below McNary Dam (Washington/Oregon, USA), natural- and hatchery-origin individuals from the Snake River (Hagerman, Idaho), natural-origin individuals from Kootenai River (near Bonners Ferry, Idaho, USA), hatchery broodstock from the lower Fraser River population (British Columbia, Canada), and hatchery broodstock from the Sacramento River population (California, USA). Sex from these individuals was identified through abdominal biopsy and/or through hatchery cross records. Markers were optimized for inclusion into the GT-seq panel (Supplemental File 1), sequenced on an Illumina NextSeq 2000 (1×100bp), and genotyped with a custom pipeline (Delomas et al., 2021).

## Results

Estimated coverage depth of the sequencing data used to construct the genome assembly depended on which published genome size (5.1pg vs. 9.6pg; Blacklidge & Bidwell, 1993; Hinegardner, 1976) was used for calculation (Table 1). The Genomescope estimated haploid genome size of the individual used for sequencing ranged from 1.514 Gbp to 1.622 Gbp, depending on assumed ploidy (2N or 4N) and kmer length utilized (Supplemental Table 1), indicating published genome sizes for white sturgeon reflect tetraploidy or hexaploidy.

Consistent with this, Smudgeplot analysis of heterozygous kmers (hetmers) indicated that the genome of the sequenced individual largely reflected tetraploidy, with an equal or smaller portion reflective of octoploidy with smaller or larger kmer lengths, respectively (Figure 1, Supplemental Figure 1). The signal of tetraploidy at longer lengths of k may reflect the products of a more derived duplication event within Acipenseriformes, while shorter values of k appear to capture sequence matches resulting from an older duplication within the sturgeon stem lineage.

**Figure 1.**
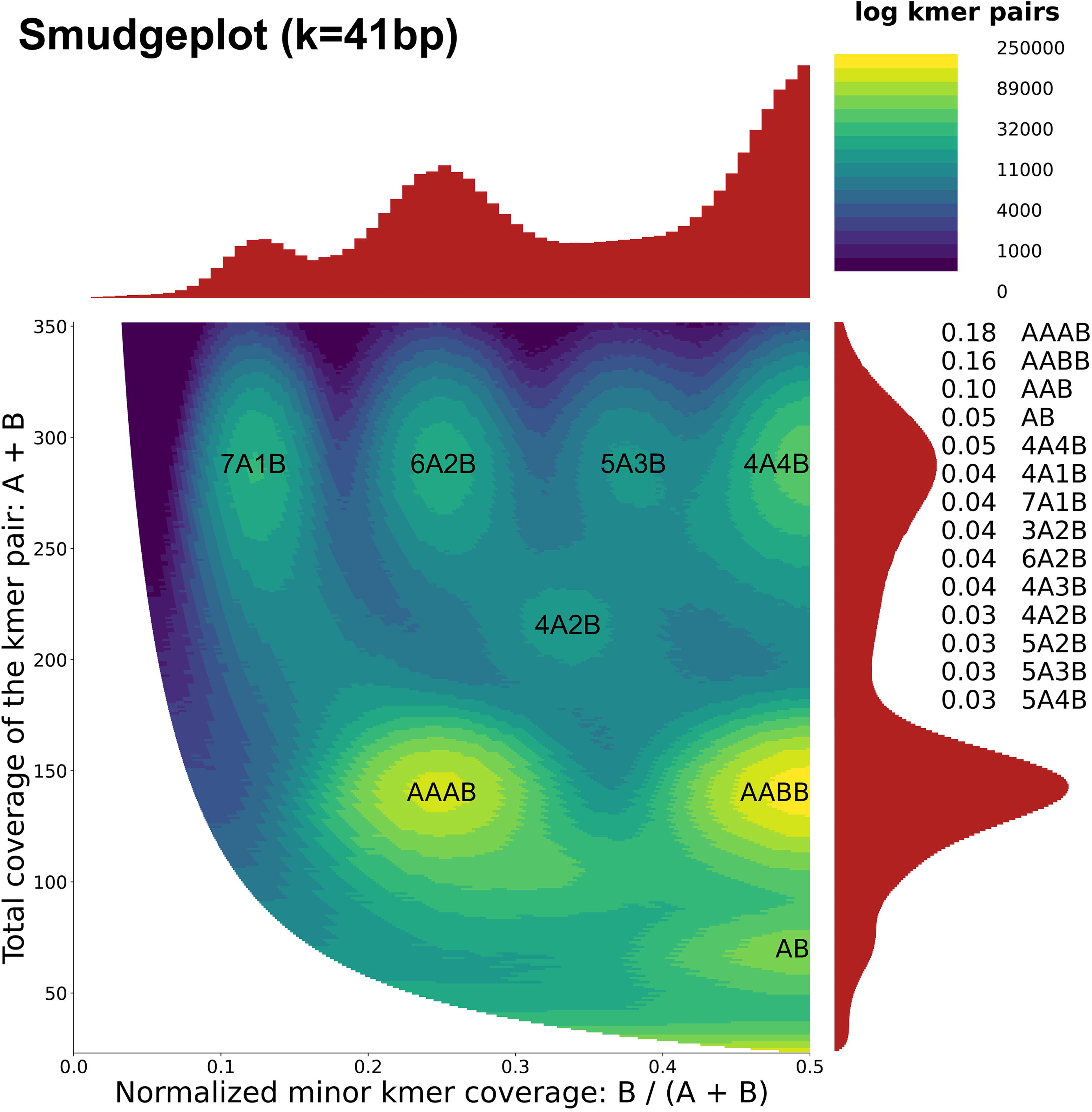
Genome content estimated from Smudgeplot analysis of heterozygous kmers with k=41bp.

The total size of the assembled contigs represented 5.96Gbp of genomic sequence, and these were assembled into initial scaffolds reflecting 6.26Gbp of genomic sequence with 4.86% gaps (Table 2). An additional 826Kbp of sequence data could not be incorporated into the assembly, which likely represents complex regions such as those with high GC content or highly repetitive sequences, as well as possible microbial contamination. Moreover, the scaffolded assembly, despite the robust structural sequence information, was notably fragmented, with an N50 of only 339Kbp for 3,870 sequences. Similarly, the largest scaffold, only 8.59Mbp, was small relative to the sterlet RefSeq assembly, which contained 52 chromosome-level scaffolds larger than this. In contrast, assessments of genome completeness based on orthologous gene content using Compleasm indicated the white sturgeon assembly was only marginally less complete than the female sterlet RefSeq assembly, with 6.1% of orthologs missing vs. 3.3% in sterlet (Table 3). Notably, both assemblies exhibited similar proportions of single copy (22.8% and 27.0%, respectively) and duplicated (67.7% and 68.9%, respectively) orthologs, despite the higher ploidy of white sturgeon.

**Table 2.** Statistics for the white sturgeon genome assemblies.

|  | **Contigs** | **Scaffolds** | **Linked Scaffolds** |
| --- | --- | --- | --- |
| Total contigs/scaffolds | 1,347,142 | 835,784 | 832,145 |
| Assembly size (bp) | 5,960,370,858 | 6,264,925,766 | 6,265,289,666 |
| Gaps size (bp) | N/A | 304,554,908 | 304,918,808 |
| Gaps (%) | N/A | 4.861 | 4.867 |
| N50 (bp) | 13,367 | 339,349 | 415,779 |
| N50 (#seq) | 90,723 | 3,870 | 1,976 |
| N90 (bp) | 1,565 | 2,015 | 2,016 |
| N90 (#seq) | 761,174 | 402,904 | 399,337 |
| Largest contig/scaffold (bp) | 359,327 | 8,590,638 | 55,262,319 |

**Table 3.** Results from Compleasm evaluation for otholog content using Actinopterygii_od12 of sterlet and white sturgeon genome assemblies.

|  | **sterlet RefSeq**<br>**(GCF_902713425.1)** | **white sturgeon**<br>**phased scaffolds** |
| --- | --- | --- |
| Complete, single copy | 27.00% (1946) | 22.80% (1643) |
| Complete, duplicated | 68.91% (4966) | 67.67% (4877) |
| Fragmented | 0.82% (59) | 3.42% (247) |
| Missing | 3.27% (236) | 6.11% (440) |

The mapping families used for scaffolding varied with respect to the number of individuals passing quality thresholds, numbers of markers captured, and the estimated recombination rates (Table 4). The map based on haploid RAD-seq data was considered to be the most accurate because it yielded a number of linkage groups that is close to the expected number of chromosomes based on karyotypic analyses (1n ∼ 135; Van Eenennaam et al., 1998), with an average rate of recombination of 0.5 per LG per meiosis. The RNA-seq-based linkage map appeared to overestimate recombination rates, possibly due to lower genotyping accuracy. Based on the anticipated and realized properties of these maps, we preferentially used the haploid map for scaffolding, supplementing with the diploid and RNA-seq-based maps where they did not conflict with the haploid map.

**Table 4.** Statistics for the linkage maps constructed from RAD-seq (Fraser River family) and RNA-seq (Snake River family) data.

|  | 'haploid'<br>(RAD) | 'diploid'<br>(RAD) | RNA |
| --- | --- | --- | --- |
| N individuals | 113 | 37 | 92 |
| total LGs | 168 | 156 | 435 |
| total cM | 9651.818 | 3815.993 | 186319.269 |
| total variants | 1678 | 951 | 11359 |

Joining of the initial scaffolds using mapped linkage group markers increased the size of the largest scaffolds in the assembly, with the largest scaffold becoming 55.2Mbp and an N50 of 416Kbp for 1,976 scaffolds (Table 2). The largest scaffolds of the final, linked assembly showed clear homology to chromosome-level scaffolds of the sterlet sturgeon, including numerous 1:2 and 1:many relationships (Figure 2). Notably, several white sturgeon scaffolds showed contiguous homology to large regions of sterlet chromosomes, with a few breaks that either reflect fission/fusion events or assembly fragmentation (e.g. white sturgeon scaffolds 1 and 3 to sterlet chromosome 3). Other scaffolds show lower degrees of linear conservation (e.g. homologs of sterlet chr. 1 and 2), reflecting higher rates of intrachromosomal rearrangement, potential non-diploid segregation, or errors in scaffolding. Similar patterns of conservation and synteny disruption are also observed between divergent paralogous chromosomes within the sterlet genome, suggesting that patterns of homology observed between white sturgeon and sterlet might reflect true differences in white sturgeon chromosomal structure. For example, sterlet chr. 7 has segments that are paralogous to chr. 13 and chr. 14, whereas each ancestral sequence corresponds to two large scaffolds in sturgeon (scaf. 7, 10, 11 and 16), indicating that sterlet chr. 7 is likely the product of a derived fission event. Overall, most sterlet chromosomes showed similar patterns of homology to two white sturgeon scaffolds, reflecting the derived paralogy from sublineage-specific whole genome duplications (e.g. white sturgeon scaf. 1 and 5 to sterlet chr. 3 in Figure 2; see also Supplemental Figure 2). Closer examination of LD patterns showed that these paralogous white sturgeon chromosomes appear to segregate independently, with a few notable exceptions (Figure 3).

**Figure 2.**
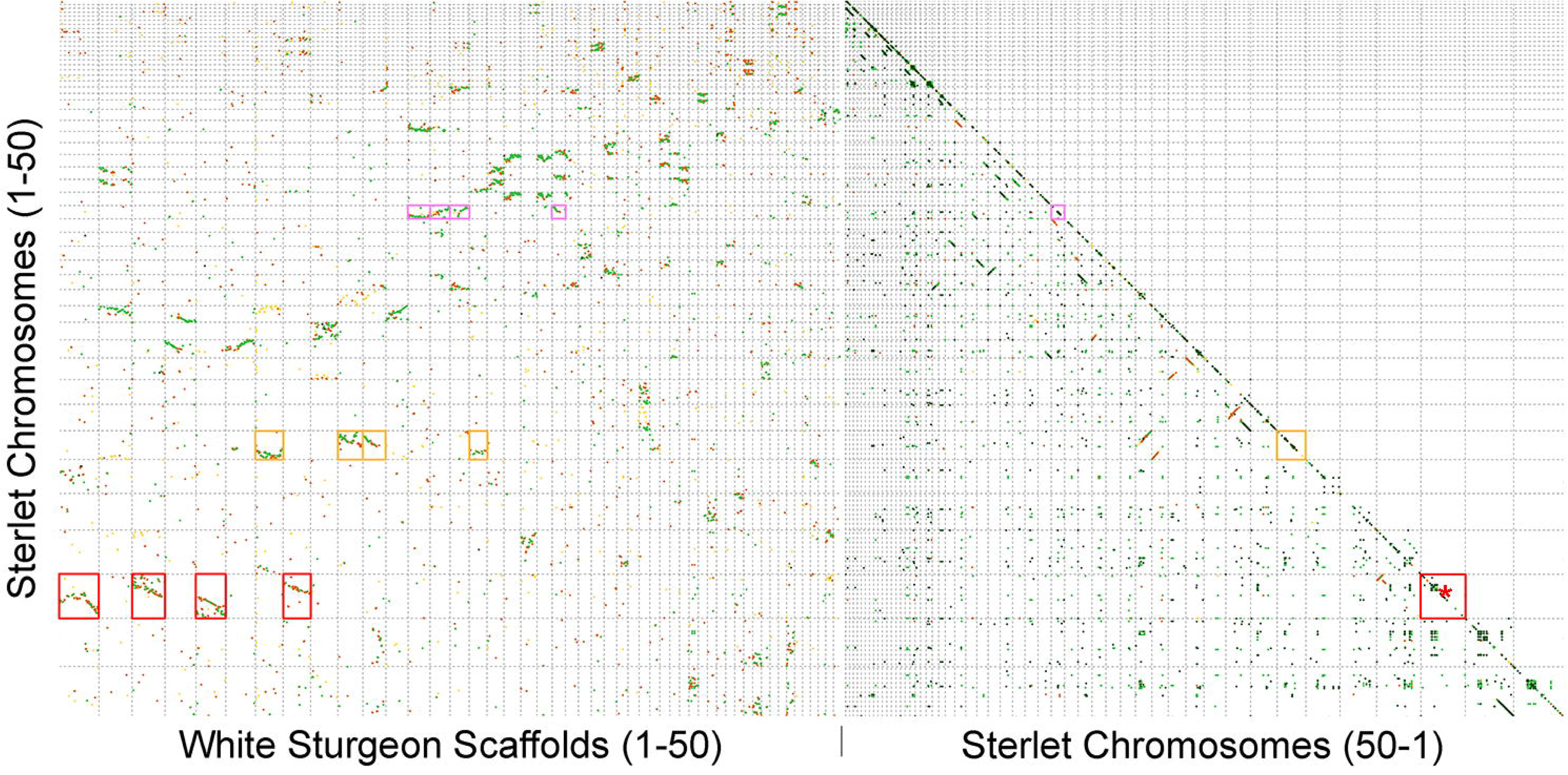
Alignment of white sturgeon scaffolds to the sterlet genome. Plots reflect alignments between the largest 50 white sturgeon and sterlet scaffolds (left). Three sets of sterlet homologs are outlined in red, orange and pink correspond to outlined chromosomes in Figure 3. Sterlet self-alignments (right) are shown to highlight older paralogy derived from a common ancestral duplication in the sturgeon lineage. A red asterisk marks the approximate location of the sterlet sex determining region.

**Figure 3.**
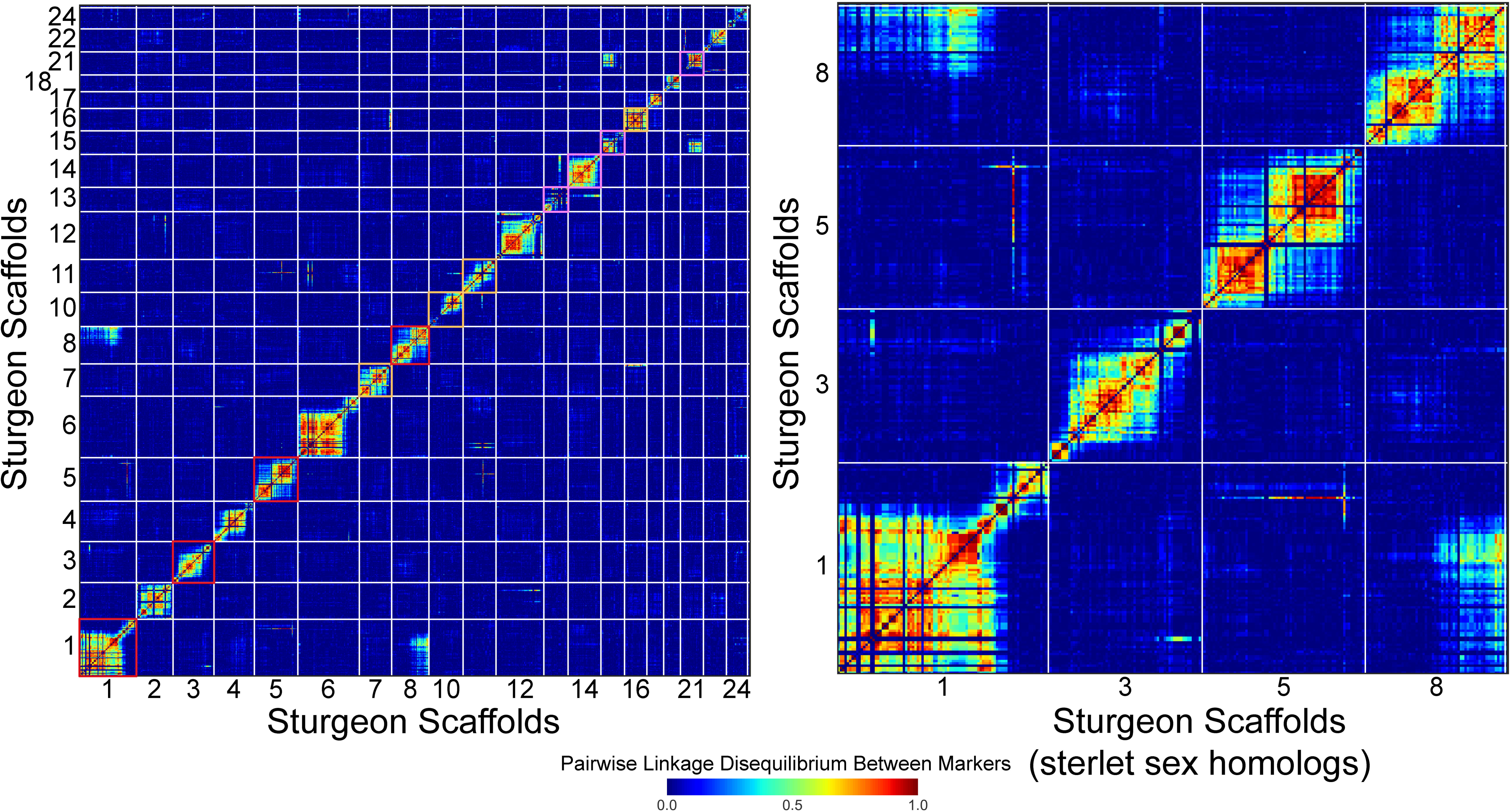
Linkage disequilibrium (ld) among segregating maternal polymorphisms from haploid segregants. A) Patterns of ld within and between the largest 25 scaffolds with segregating maternal markers. Chromosomes outlined in red, orange and pink correspond to outlined chromosomes in Figure 1. B) Patterns of ld within and between four scaffolds that are homologous to the sterlet sex chromosome.

The white sturgeon scaffolds identified through *blastn* searches with the sterlet sex-specific regions showed contiguous homology to those sterlet extractions, including in the respective sex-specific regions (Supplemental Figure 3). Notably, the regions corresponding to the sterlet male extraction (putative-Z) were joined into the final white sturgeon scaffolds 1 and 5, while the much shorter female-corresponding scaffold (putative W), only 108Kbp, was not incorporated into the linkage map, exhibiting segregations patterns that were not correlated with portions of the genome that were captured by linkage analyses. As noted above, these two final white sturgeon scaffolds are part of a group which form contiguous and duplicated homology with sterlet chromosome 3, wherein the sex-specific regions reside. Notably, these two joined scaffolds show evidence of latent linkage disequilibrium (Figure 3), suggesting the possibility that they continue to pair at meiosis and exhibit ongoing recombination.

The two amplicons developed from these homologous sex-specific regions showed strict linkage in genotyping, with only individuals exhibiting the putative-W-specific SNP haplotype also exhibiting successful amplification of the putative-W-specific presence/absence marker positioned in the adjacent, unalignable region. Also notable, the putative-W-specific haplotype, when present in individuals with >80% overall genotyping success, reflected approximately 1/4th of the sequenced reads for that locus (mean 0.251, stdev 0.023). The overall efficacy of the presence or absence of the putative-W-specific haplotype or amplicon at predicting phenotypic females and males, respectively, was high (96.6% and 81.5%, respectively), providing a strong overall association with sex (□^2^ p-value < 2.7×10^-37^). However, the efficacy varied across stocks (Table 5): for individuals in the lower Columbia River, whose sex was confirmed both by abdominal biopsy and reproductive success (parentage), accuracy was 100% in both sexes, but for males in the Sacramento Basin, presence of the putative-W-specific markers was higher than expected (66%). Other stocks showed intermediate accuracy for one or both sexes.

**Table 5.**
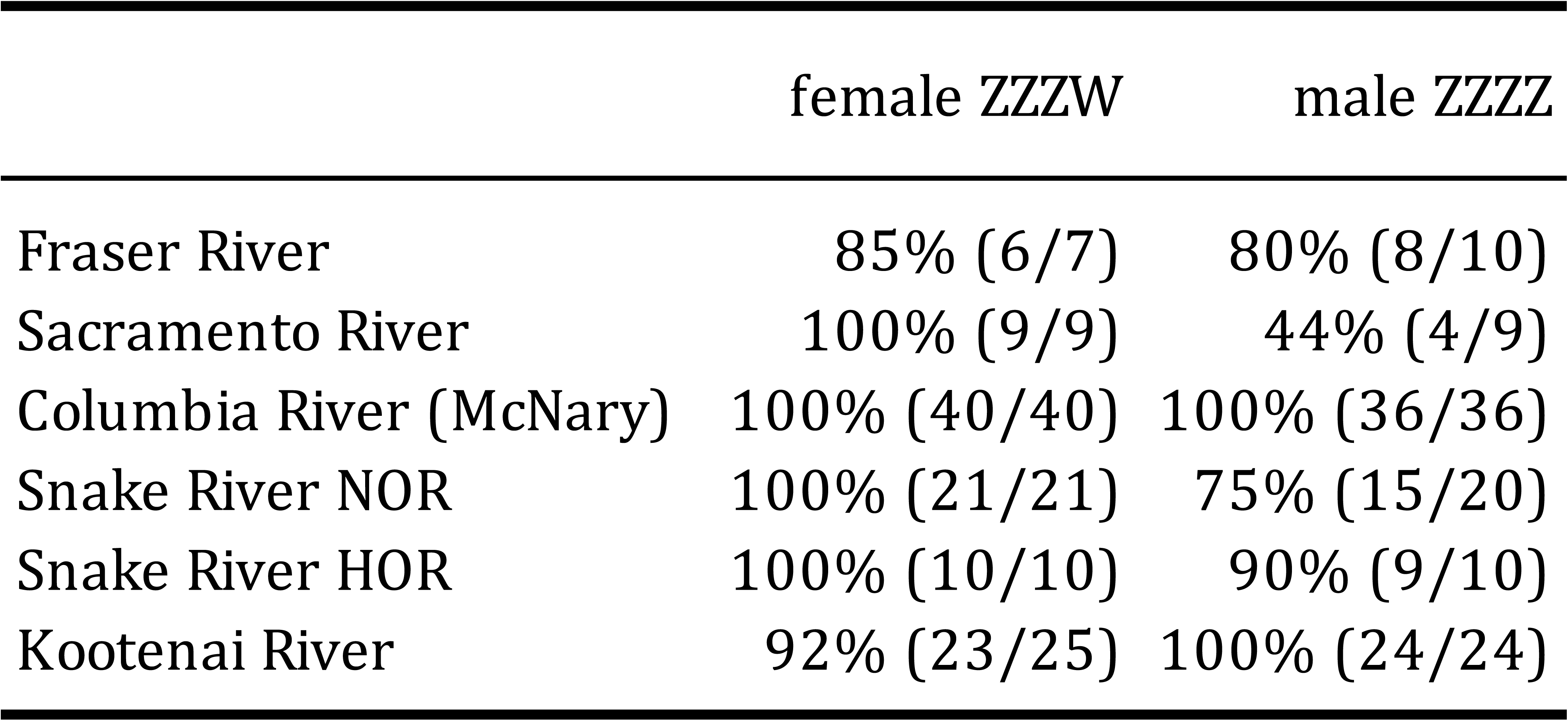
Proportions of known sex white sturgeon exhibiting the predicted genotypes for molecular markers for phenotypic sex. In parentheses, correctly predicted and all samples for a given sex. For the Snake River, natural-origin (NOR) and hatchery-origin (HOR) individuals are distinguished.

## Discussion

This study provides a novel genomic resource for white sturgeon which will aid in the conservation of this charismatic and imperiled species as well as help facilitate research into the process of genome evolution for polyploid vertebrates. However, this resource remains in a notably fragmented state compared with the sterlet assembly, which partially reflects the higher ploidy and inherently higher paralogy within the white sturgeon genome. We utilized a pipeline that has been successfully applied previously for species with autopolyploid genomes, but it could well be that the complex nature of the white sturgeon genome, with compounded autopolyploid events, will require the use more advanced sequencing approaches such as long-read and Hi-C libraries. Despite the residual fragmentation in the assembly provided here, orthology assessments indicate that the current assembly contains most of the gene content of other sturgeon genomes. Thus, the present white sturgeon genome assembly will be useful for a variety of genome-dependent genetic studies.

To demonstrate the utility of the present genome assembly, we have utilized it in pursuit of genetic markers for phenotypic sex in white sturgeon. White sturgeon are widely used in aquaculture for both meat and caviar production, with the latter relying on the production of mature, female fish. Having molecular markers to predict the sex of fish prior to maturity would assist commercial operations in managing limited resources most effectively. Further, conservation aquaculture programs have been implemented to bolster declining populations or limited recruitment for a number of white sturgeon populations (Mallette, 2014). These programs are at times forced to selectively cull slower growing young in order to maximize resource efficiency but without knowing the impact of these events on residual sex ratios. Moreover, while hatchery fertilization procedures have been optimized in order to minimize the incidence of spontaneous autopolyploidy in hatchery offspring, selective culling for higher ploidy continues to be necessary but without knowledge of the dynamics of sex among autopolyploids.

Within the present white sturgeon genome assembly, we identified regions which show strong homology to the sex-specific regions of other sturgeon (Kuhl et al., 2021). Moreover, markers developed for two sequences within this region showed strong but variable association with sex. While the listed primers for these markers have been optimized for inclusion in a multiplex, GT-seq panel, they could be adapted to be used as a standalone qPCR or visual assay (e.g., Kanefsky et al., 2022), e.g. presence/absence of the female-specific marker, *p455160ex*, possibly with the shared marker, *zwhr*, as a positive control (in which case the primers of one marker should be 5’ elongated to distinguish amplicons based on size). PCR efficiency or primer concentrations notwithstanding, the expected amplicon concentration for females would be ∼1:4 *p455160ex*:*zwhr*.

However, like any new technology, we urge caution in implementing this assay for phenotypic sex and suggest validating the specific association with sex using known sex fish in individual white sturgeon stocks. As indicated by our tests, while the association with sex for these markers is strong, it varies significantly among the stocks tested here. There are several explanations for variable association. First, while the association of these sex-specific, unalignable regions appears clear for white, sterlet, and other sturgeons (Degani et al., 2022; Kuhl et al., 2021; Scribner & Kanefsky, 2021; this study) it may only be linked to variants that contribute to rather than being directly involved in sex determination *per se*. The RefSeq annotation for the female sterlet assembly indicates that the sex-specific region resides in the first intron of the *ofcc1* gene (“orofacial cleft 1 candidate 1”; LOC117394325), a transcribed but non-coding region with a possible regulatory role. While this gene is largely uncharacterized in fishes and has not been directly proposed as a sex determining gene, we note that it is strongly expressed in embryos as well as the testis and ovaries of male and female zebrafish (*Danio rerio*) (expression for ENSDARG00000104791 on Bgee, https://www.bgee.org). Whether this gene, and the sex-associated region within it, is causal or simply linked to sex-determining variation remains to be determined. Second, patterns of residual linkage suggest that two scaffolds containing sequences homologous to the sterlet male-specific region (putative Z chromosomes) experience some level of ongoing recombination. In many species, recombination is suppressed between the sex chromosomes, or sex regions of pseudo-autosomes. Thus, it remains to be resolved how distantly situated from the actual locus or variant(s) of determination these markers lie and how much even low levels of recombination may diminish association in any given stock. Third, while most species with genetic sex determination exhibit a single, ‘master’ sex-determining gene, sex determination could be polygenic in sturgeon. Indeed, Degani et al. (2022) identified a number of sex-associated SNP variants in the Russian sturgeon (*A. gueldenstaedtii*), a distantly related, octoploid species. It is possible that there are additional, unlinked variants which play a larger role in mediating sex determination in some stocks of white sturgeon. Finally, although it has been accepted for some time that sturgeon exhibit a female-heterogametic sex determination system (ZW), it is not clear that sex determination is fully genetic in white or other sturgeon. If not, environmental conditions that vary among stocks, or between hatchery and natural-origin fish, could diminish the association of molecular markers for sex despite strong genetic linkage.

Beyond the use of additional sequencing technologies to further resolve the polyploid white sturgeon assembly, greater utility of a white sturgeon genome resource will come from a ‘pan-genome’ of several assemblies that represents a compendium of sexual, population genetic, and other relevant variation from across a species (e.g., Gong et al., 2023). While this term was originally applied to microbial species which exhibit a ‘core’ set of genes present across all individuals and a large library of ‘accessory’ genes, it has been increasingly applied to vertebrate species as the unexpected degree of genomic variation among individuals has become apparent. As has been shown for species of salmonids, gene content and structural variation among individuals can play a critical role in mediating major life history variations relevant for both aquaculture and population management (e.g. Pearse 2019, Bertolotti 2020). For an ancestrally octoploid and functionally tetraploid species with strong genetic structure among some stocks, the potential for important structural variation to segregate is great, but beyond even knowing of its existence, representing and making use of that variation requires a genomic resource that can reflect genomic variation of multidimensional complexity.

Representation and bioinformatic tools making use of pan-genomes is an active area of research, but the complexity reflected in a single white sturgeon genome assembly highlights the opportunities created by progress in this area.

## Data Availability

Raw data for linkage map creation are available from the NCBI Short Read Archive (PRJNA1271479). The final assembly and raw data used to construct it are available from NCBI Genome and Short Read Archive, respectively (PRJNA1358253).

## Supporting information

SFile 1 GTseq panel

## Acknowledgements

The authors acknowledge Ben Sutherland and Dan Baker, and Gary and Linda Lemmon, who provided the Fraser River and Snake River white sturgeon families used to construct linkage maps. Samples from the focal Snake River white sturgeon were collected by Nate Campbell and Andrew Matala. Laboratory staff at the Hagerman Genetics Lab, in particular Stephanie Harmon, Kim Vertacnik, and Rachael Kane, facilitated library construction and sequencing of the linkage map data.

## Supplemental Files

Supplemental File 1. GT-seq panel for white sturgeon, revised Sept. 2025

---

**Supplemental Figure 1.**
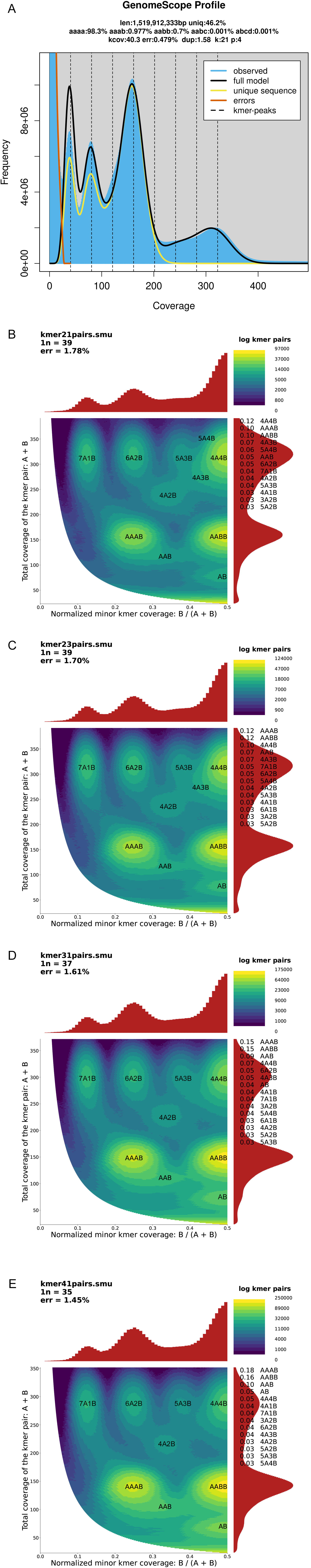
Genome content analysis using kmers. A) Representative Genomescope results, with k=21bp and ploidy=4N. B) Smudgeplot analysis with k=21bp. C) Smudgeplot analysis with k=23bp. D) Smudgeplot analysis with k=31bp. E) Smudgeplot analysis with k=41bp.

**Supplemental Figure 2.**
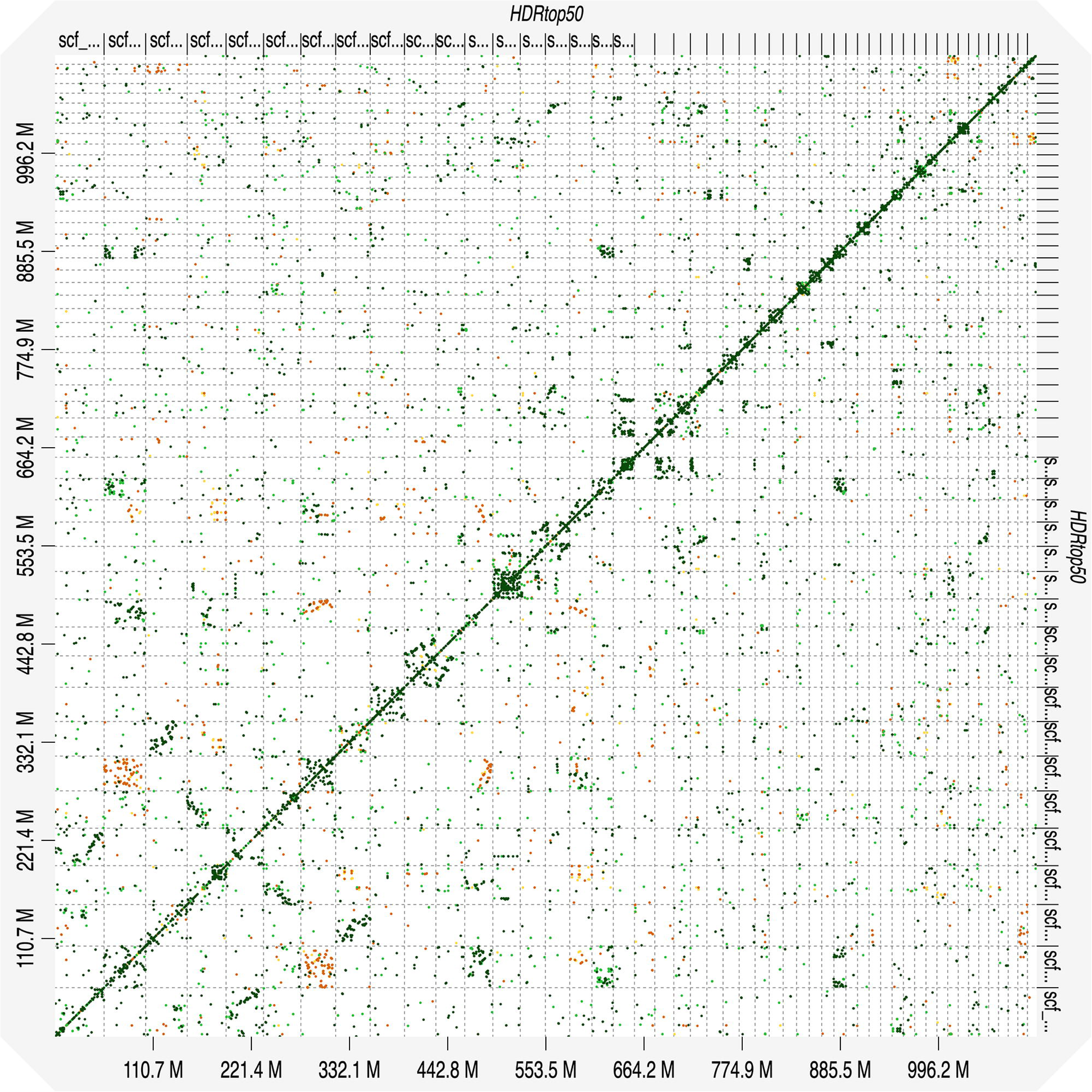
Alignment of white sturgeon largest 50 scaffolds to themselves. Scaffold ordering follows Figure 2.

**Supplemental Figure 3.**
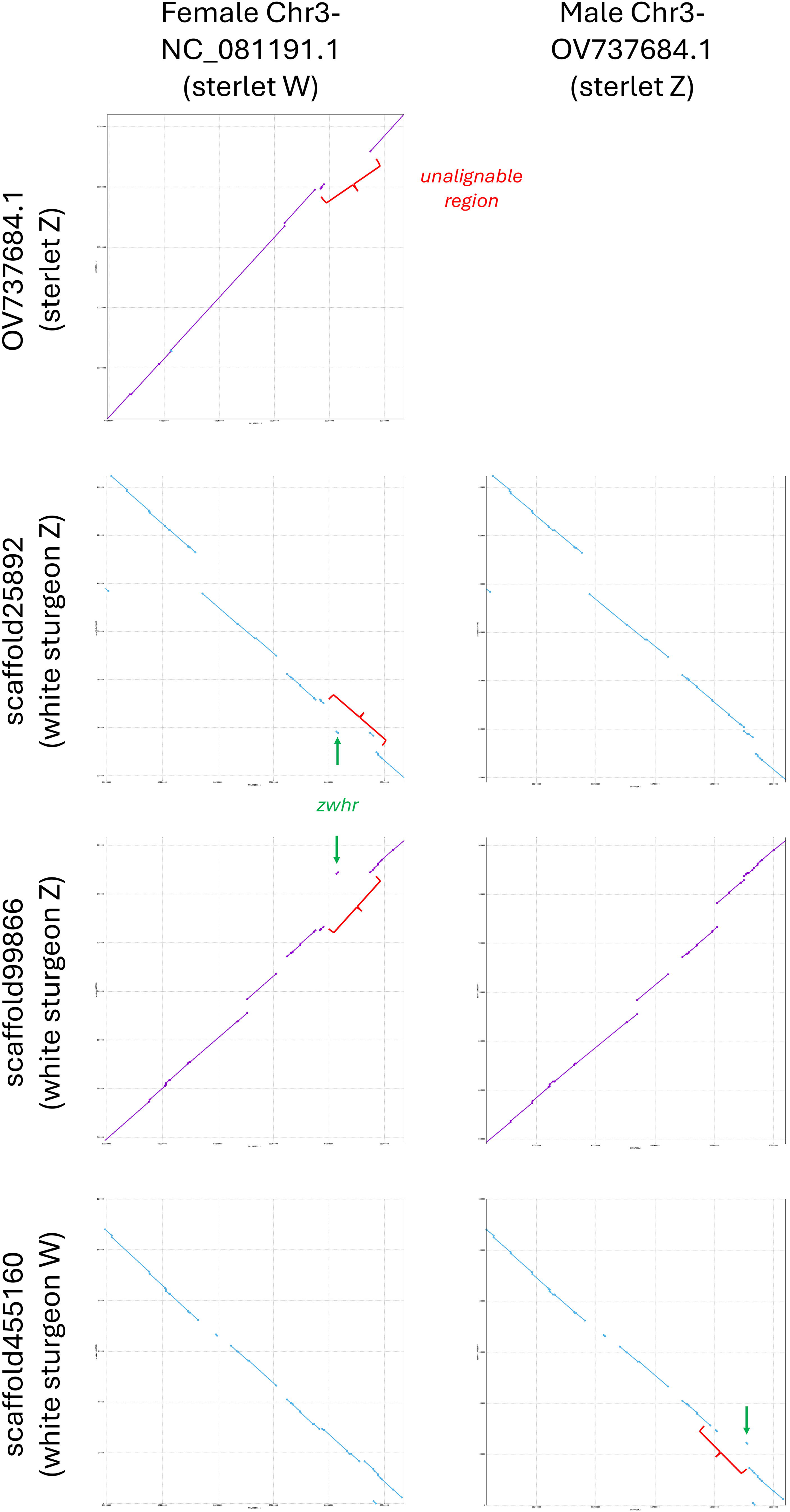
Mummer alignments of the sex-specific and flanking regions of sterlet sturgeon (from Du et al. 20XX) and three homologous scaffolds from the white sturgeon phased assembly.

**Supplemental Table 1.** Haploid genome sizes estimated with Genomescope from various kmer lengths and diploid or tetraploid status.

| **kmer** | **ploidy** | **min (bp)** | **max (bp)** |
| --- | --- | --- | --- |
| 21 | 2N | 1,514,830,630 | 1,519,678,665 |
| 21 | 4N | 1,515,246,697 | 1,519,912,333 |
| 23 | 2N | 1,530,386,796 | 1,535,475,021 |
| 23 | 4N | 1,532,477,521 | 1,536,954,560 |
| 31 | 2N | 1,574,211,122 | 1,580,387,873 |
| 31 | 4N | 1,582,577,639 | 1,586,632,609 |
| 41 | 2N | 1,604,596,808 | 1,612,141,825 |
| 41 | 4N | 1,618,902,487 | 1,622,808,454 |

